# Larval zebrafish exhibit collective motion behaviors in constrained spaces

**DOI:** 10.1101/2021.07.09.451841

**Authors:** Haider Zaki, Enkeleida Lushi, Kristen E. Severi

## Abstract

Collective behavior may be elicited or can spontaneously emerge by a combination of interactions with the physical environment and conspecifics moving within that environment. To investigate the relative contributions of these factors in a small millimeter-scale swimming organism, we observed larval zebrafish, interacting at varying densities under circular confinement. Our aim was to understand the biological and physical mechanisms acting on these larvae as they swim together inside circular confinements. If left undisturbed, larval zebrafish swim intermittently in a burst and coast manner and are socially independent at this developmental stage, before shoaling behavioral onset. We report here our analysis of a new observation for this well-studied species: in circular confinement and at sufficiently high densities, the larvae collectively circle rapidly alongside the boundary. This is a new physical example of self-organization of mesoscale living active matter driven by boundaries and environment geometry. We believe this is a step forward toward using a prominent biological model system in a new interdisciplinary context to advance knowledge of the physics of social interactions.

## INTRODUCTION

The emergence of complex collective behavior of natural and artificial motile agents has long been a question of interest to scientists in many disciplines. The transition from disordered to ordered collective motion can be seen across scales, from micron long bacteria and colloids, to millimeter long ants and bees, to centimeter long crickets and bristle-bots, and even to meter-long fish and humans (1–3). The complex behavior of flocks of birds, colonies of ants, swarms of bees and schools of fish emerges from the interactions of the constituent parts of the respective systems. While similarities in the patterns that such groups produce have suggested general principles governing the self-organization (4), it is also becoming clear that the specific patterns depend on the type of motile agent, scale, and also the type of interaction. For example, for fluid-immersed micro-scale units such as motile bacteria and colloids, it has become clear that mechanical interactions often mediated through the liquid are paramount to the type of eventual patterns (5,6). For larger animals such as birds, mechanics are not the only factor as others may become more prominent, e.g., visual input for birds (7), environmental factors for bees (8), sensory stimuli or social cues for humans (9).

Despite the large effort in studying the emergence of collective motion for various motile agents, little has been done to study how this behavior changes when the agents are confined, whether by hard walls or soft impediments. Recent work has shown that when swimming bacteria, colloids, spermatozoa, and even bristle-bots are placed in circular or racetrack dishes, then they will spontaneously start to circulate (5,6,10–13). Even soft confinement can lead to locust milling (4) and human mosh pits (14). Our approach was motivated by the need to develop a model where behavior can be observed easily, which is amenable to neurobiological perturbations, and which generates interesting and quantifiable individual and collective behavior. The popular biological model organism *Danio rerio*, the zebrafish, has been a top-tier model in neuroscience over the recent decades, and is relatively simple to observe. Since this millimeter-scale swimmer has a non-trivial behavior (i.e. moves discontinuously and can change direction suddenly) influenced by fluid mechanics as well as by sensory stimuli (15–18) we expected to see a rich and complex collective motion sharing similarities yet different from other previously studied motile agents. Adult zebrafish have been studied extensively in both individual and collective contexts (19–24). However, at the 5 days post-fertilization larval life stage, when zebrafish are approximately 4 mm in length, before the onset of shoaling behavior (25,26), they utilize different movement patterns from adult fish. Larval zebrafish swim in what is often termed a beat-and-glide or burst-and-coast discontinuous manner; they swim in bouts of movement followed by pauses (27). There has been a steady development in the literature of knowledge and tools regarding the behavioral repertoire of larval zebrafish, including work on the range of behaviors exhibited (27–32), as well as focus on specific canonical responses such as the escape response (33–36) and prey capture (37–42). Larval zebrafish make an excellent model to observe behavior and to investigate the neural circuits underlying this behavior. In some cases, components of the neural circuits have been identified down to the specific nuclei or individual neurons (28,33,43–50). The tools available in zebrafish to potentially explore the neural circuitry responsible for such behavior led us to test whether larval zebrafish respond collectively to high densities in confinement.

At this life stage, larval zebrafish placed in low density have a usual social avoidance area of approximately 50 mm^2^ surrounding their body where they prefer an absence of conspecifics and will initiate escape responses to avoid them (51). However, the set of observations we report here included a range of densities where larvae were forced to interact with conspecifics and the arena walls which in some cases did not permit them to maintain their preferred social avoidance area. We observed that when in confined environments at sufficiently high densities, larval zebrafish may spontaneously collectively perform a novel circling behavior along the confining dish. Here we report conditions under which collective circling behavior may be elicited in larval zebrafish, a tractable model system, opening new avenues of investigation.

## MATERIALS AND METHODS

### Animals

Larval zebrafish used in these experiments were 5 days post-fertilization (dpf) AB wild-type (origin: ZIRC stock center, Eugene, Oregon) reared in an incubator at 28.5°C with a 14L:10D light cycle. Larvae were generated from an adult colony maintained at NJIT in the Severi lab under Rutgers University-Newark IACUC oversight, PROTO201800041. The same larvae were used for each trial. Videos were captured approximately 1-5 minutes after larvae were placed in the behavior enclosure to allow time for acclimation following handling.

### Acquisition

High-speed videos were collected on a custom-built setup (**Figure 1A, Supplemental Table 1**). A high-speed camera (Mikrotron GmbH, Germany) attached to a rail (ThorLabs) fitted with a 35 mm F1.4 lens (Fujinon) and an 850 nm bandpass filter (Midwest Optical) were used to acquire images to a Dell Precision 5820 computer fitted with a frame grabber (National Instruments) and running custom-written LabView software (National Instruments, available upon request) saving TIFF image stacks for each trial. Larvae were illuminated with 850 nm IR LEDs which are not within their visible spectra (Waveform Lighting) under an acrylic platform stabilized by Thorlabs components covered with light diffusers (Pro Gel, B&H Photo) within a custom-built enclosure (MiniTec Framing Systems, LLC) which was left open to room light. Videos were acquired at 200 Hz with 1423 μs shutter speed at 648 x 648 pixel resolution, and trials were 6000 frames or 30 seconds in duration. Animals were recorded at room temperature during daytime in round petri dishes with 5.4 cm diameter.

**Figure 1.**
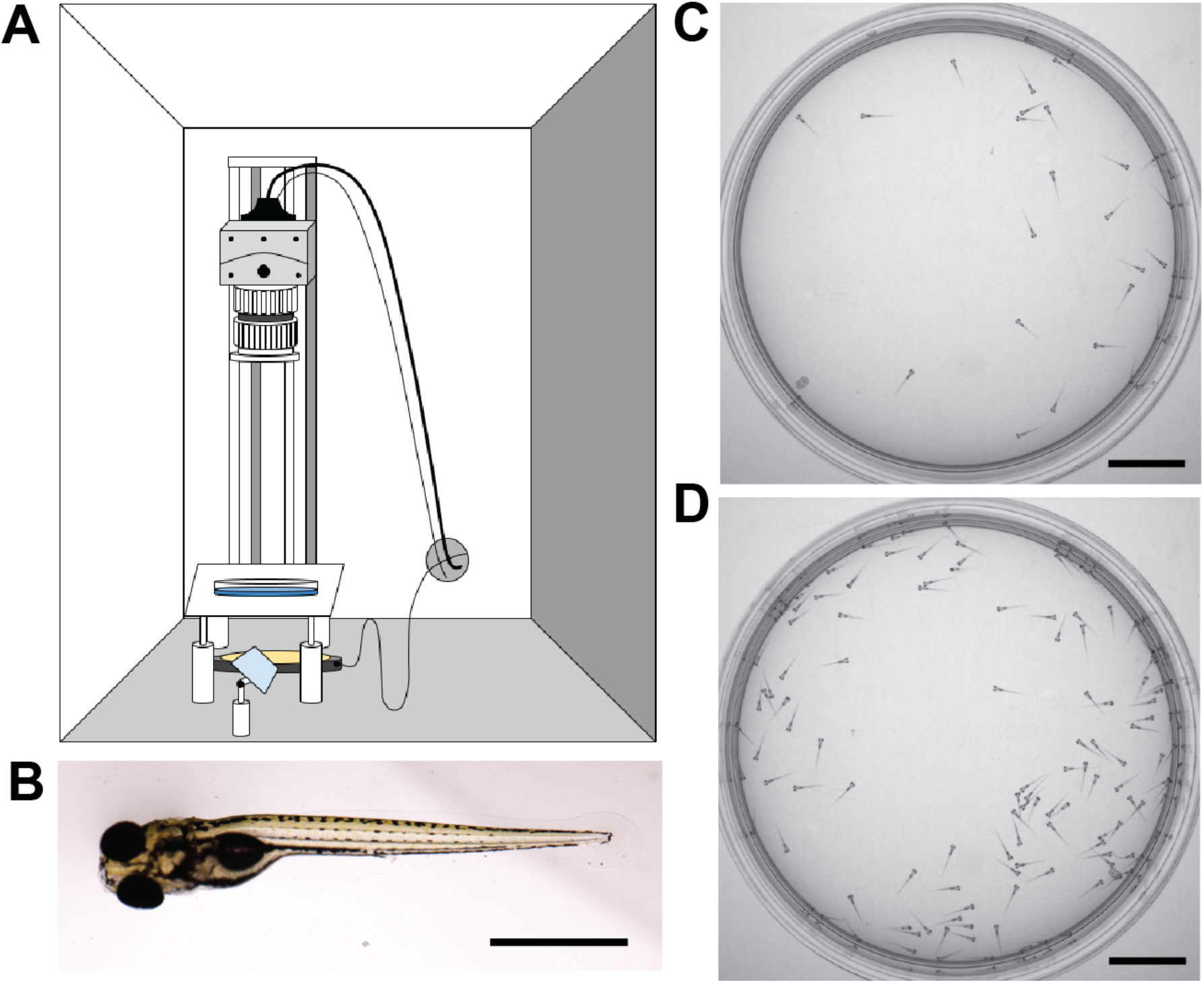
Experimental observation of larval zebrafish swimming collectively at various densities. **A**. Schematic of experimental set-up. **B**. Image of a larval zebrafish at 6 days post-fertilization, scale bar 1 mm. Rostral (head) is to the left. **C**. 30 larval zebrafish in the arena as captured by the camera. **D**. 130 larval zebrafish in the arena. Scale bar for C-D 1 cm.

### Analysis

To generate tracks from raw videos, we utilized trackR, an R package written by Dr. Simon Garnier (52). trackR is an object tracker for R allowing users to perform multi-object video tracking by background subtraction and adaptive thresholding. This tracking solution allows defining parameters of object detection, and distinctions in object continuity over the course of input videos. Together with an understanding of the optimal video acquisition pipeline and ideal capture specifications, trackR outputted well-tracked object traces as .csv files. These were imported to MATLAB and plotted. To determine how position varied with radial distance using FIJI (NIH) and MATLAB, videos were reduced by ¼ in FIJI and cropped to exclude the pixels outside the arena. A standard deviation z-projection was applied to the image stacks to create a single image using FIJI. This process outputs a single image where each pixel represents the standard deviation value over all images in the stack at that particular pixel location (https://imagejdocu.tudor.lu/gui/image/stacks#zproject). Using the MATLAB function average_radial_profile_2 (Image Analyst, Mathworks author id:31862) the average radial profile was calculated from a center location of each image and plotted.

For determination of the circling behavior (**Table 1**), a pair of qualitative assessments of the captured videos were used. First, when watching the videos at 30 Hz playback and paying attention to the region just inside the arena boundary, some collective circling instances were immediately obvious based on easily discernible rotational movement. When many conspecifics began moving in a coordinated manner around the edge of the dish in a single major direction, we took this to be circling behavior. A second qualitative identifier of circling motion was a correlate of the behavior that arises due to fluid flow. When a significant number of the larvae are circling near the boundary of the dish, a radial region of the fish within the circling ring will exhibit larvae in counter-rotation, moving in the opposite circular direction to the major circling direction of the outer ring. This is characterized through the obvious lack of self-propelled movement (the larvae themselves are static), and oftentimes a drifting motion in a backward direction which is not a gait present in this species. Based on our understanding of fluid motion (6), this is strongly indicative of coordinated circling in one major direction. With either or both of these qualifications met, we could categorize a captured video to have circling behavior.

**Table 1:** Table of density across trial noting when circling behavior was observed.

| **Arena Diameter (cm)** | **Fish Quantity (n)** | **Density (fish/cm<sup>2</sup>)** | **Circling Behavior** | **Trials** |
| --- | --- | --- | --- | --- |
| 5.4 | 5 | 0.218 | not observed | 5 |
| 5.4 | 10 | 0.437 | not observed | 5 |
| 5.4 | 30 | 1.310 | observed 1/5 | 5 |
| 5.4 | 50 | 2.183 | observed 2/5 | 5 |
| 5.4 | 70 | 3.057 | not observed | 5 |
| 5.4 | 100 | 4.367 | observed 3/6 | 6 |
| 5.4 | 130 | 5.677 | observed 8/10 | 10 |

To determine the distance traveled over tracks in separate regions within the dish, tracks were imported to MATLAB and the arena boundary and center were determined using a custom function, with a radius of 310 pixels. The inner circle was 70% of the radius (217 pixels) with the same center coordinates (**Figure 3A-B**). Tracks within those regions were segregated and distance across all tracks for a given trial were calculated for three trials with 30 larvae and three trials with 130 larvae over 1497 frames of the trial. Each trial and the means were plotted along with the standard error of the mean (**Figure 3C**). A standard t-test was applied between groups (ttest2 function in MATLAB) and p-values < 0.05 were considered significant.

## RESULTS

We set out to observe larval zebrafish behaving spontaneously under confinement at various densities and to determine whether collective behavior emerged. Observing larval zebrafish behaving spontaneously at varying densities, from 5 to 130 individuals, in a 5.4 cm diameter arena (**Figure 1**), we found the animals moved freely within the arena. We systematically tested a range of densities, and found that at low densities the animals behaved relatively independently of each other, though the circular wall affected their motion as they tended to swim or stop by it more often. At high densities however the animals exhibited a high-speed circling behavior, with incidence increasing alongside density. At this density the larvae can no longer ignore the presence of other larvae as well as the boundary. The circling behavior appeared to initiate on occasions when larvae came close to each other, producing a response in the contacted larvae, and that the size and shape of the arena and the interaction with the wall produced a group of larvae circling near the edge of the arena (**Figure 2, Supplemental Movie 1**).

**Figure 2.**
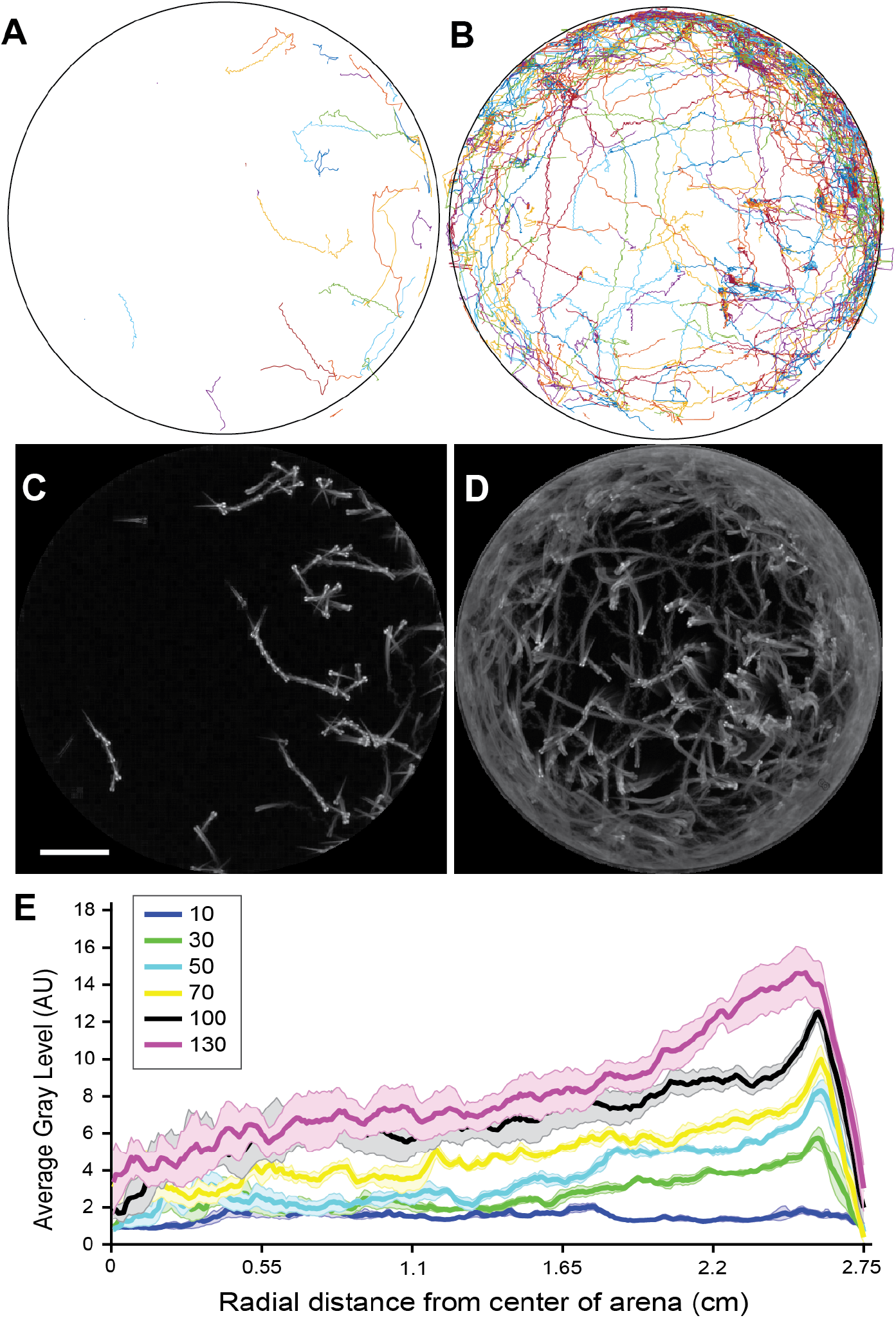
Distribution of larval positions varies as a function of distance from the arena wall. **A**. Tracked trajectories for a single trial with 30 larvae. **B**. Tracked trajectories for a single trial with 130 larvae. **C.** Standard deviation z-projection for the frames corresponding to A. **D.** Standard deviation z-projection for the frames corresponding to B. **E.** Average gray area of standard deviation z-projections plotted from the center of the arena to the border of the arena for 5 trials at each density. The mean is shown in bold and the standard error of the mean is shaded in a color corresponding to the density. Densities tested were: 10 larvae, 30 larvae, 50 larvae, 70 larvae, 100 larvae, and 130 larvae, in the same arena. Scale bar for A-D 1 cm.

Tracking the positions of the animals confirmed there was a propensity to spend time in the outer region of the arena in proximity to the arena walls (**Figure 2A-B**). While a preference for the arena edges is well noted and was found at densities of 30 or greater larvae, the distribution in spatial position was highly biased toward the outer circumference of the arena at higher densities (**Figure 2 C-E**). These higher densities of 100 larvae per 5.4 cm dish and greater correlated with more frequent observation of the circling behavior (**Table 1**). We observed instances where the collective circulation slowed down, stopped, or initiated in competing directions with one direction of flow emerging as the dominant direction. This needs to be explored in future work investigating the factors that determine initiation, stopping, and direction of motion, keeping in mind that the fish can interact with others and the boundary not just through direct contact but also through fluid-mediated mechanical forces. Indeed, when the traveled distances of each tracked position were plotted and separated into the wall-adjacent boundary region of the arena and compared to the central inner circle away from the boundary (**Figure 3A-B**), there were significant differences in the distance traveled when comparing high and low densities, and when comparing the two spatial regions at high densities (**Figure 3C**). These observations led us to conclude that larval zebrafish may be utilized as an interesting model to ask both mathematical and neurobiological questions about collective motion of swimming animals in mesoscale fluidic environments. This behavior, especially at an early developmental stage when social abilities are not yet developed, gives rise to many potential investigations unique to this species.

**Figure 3.**
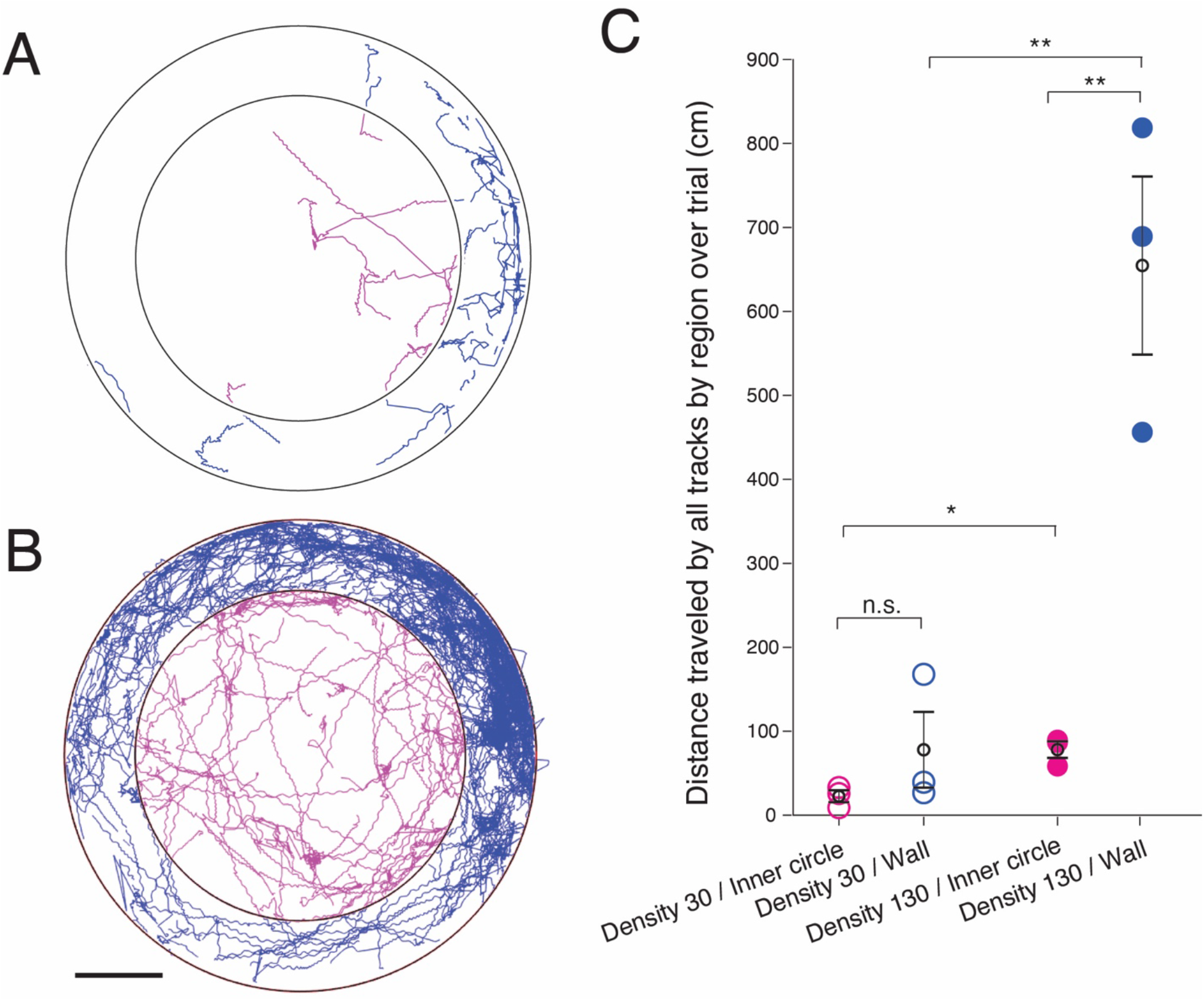
Larval zebrafish traverse space differently at different densities when bordering the arena wall in comparison to in the center of the arena. **A.** Trajectories for a trial with 30 larvae segregated into an outer ring adjacent to the wall (blue) and the arena center (magenta). **B.** Trajectories for a trial with 130 larvae segregated into an outer ring adjacent to the wall (blue) and the arena center (magenta). Scale bar for A-B 1 cm. **C.** Summed distance traveled in cm for all tracks in a trial. Trials with 30 larvae (open circles) were compared to trials of 130 larvae (filled circles) and wall-adjacent regions (blue) were compared to the inner circle of the arena (magenta). Means per group are in black open circles and error bars are the SEM. n.s. = not significant, * p <0.05, ** p < 0.01.

## DISCUSSION

The collective motion of animals and other active agents in enclosed areas is an evolving but promising area of study. We see similarities but also differences in the self-organization of larval zebrafish to other types of motile agents under circular confinement. Here we refer to collective motion simply as an emergent group behavior that only occurs as a function of the interactions of the conspecifics and their environment and would not occur if a single larva were in isolation. At high densities the larval fish transition to ordered circulation alongside the dish boundary and tend to be found swimming closer to the walls than at low densities. A largely similar circulating collective motion pattern is seen across scales from single-cell organisms to humans when the motile units are placed in hard or soft circular enclosures. And yet the specific physical and neurobiological capabilities of larval fish give rise to distinct behavior. Their non-uniform speeds may be influencing the non-uniform circulation which at times may stop or even reverse direction. Their preferred social distancing, possible visual cues, and fluidic interactions may explain why they can mostly be found at a certain distance from the confining wall that increases with density.

Unlike smaller swimmers like bacteria, algae or spermatozoa (53), these relatively larger larval fish have more complex individual motion patterns. They are well-studied however, and much is known about their biology, locomotion, individual, and social behavior. Groneberg et al., 2020 shows that the preferred distance between animals changes due to early life social interaction, and that these responses are driven by vision and by the sensory lateral line, which senses water flow around fishes (51). In future work, we hope to experimentally manipulate these various sensory inputs and systematically study the triggers for collective circling. It is highly advantageous to develop collective motion paradigms in a model system with an extensive genetic and optical toolkit to allow experimenters to observe and manipulate neural circuits (54–63). Insects are also known to display a transition from disordered to ordered movement with increasing density, most famously in locusts (4). Zebrafish sit in an advantageous space between invertebrate models which are easily studied and where sensory systems may be perturbed to investigate mechanisms, such as in *Drosophila* (64), and humans which display complex behavior yet the neurobiological underpinnings are often inaccessible, although techniques like fMRI can allow some measurement of neural activity of human individuals during complex social decision making (9).

While thigmotaxis or “wall-hugging” as a response to anxiety-inducing stimuli has been well documented in larval zebrafish (65), it is interesting to consider this emergent circling behavior in the context of social anxiety caused by crowding and confinement. It was observed in work involving small groups of zebrafish at the same life stage with much smaller arenas housing seven larvae at a time, that one larva in the group could set off chain reactions of escape responses: if one animal escaped it would collide with another setting off a domino effect (30). It’s possible this emergent collective circling results from these same chains of escapes, created by the interaction with the confinement of walls and the high density of conspecifics, which then catalyze this circling behavior. In future work we are interested in identifying mechanistic drivers of transitions between states, as has been identified in other species (66).

Another interesting observation was noted just inside the extreme edges of the arena, where immobile larvae can be seen drifting rearwards, in counter-rotation with the adjacent larvae circling at the dish circumference. We presume this can be attributed to fluid flow as the animals do not appear to be oscillating their tail or actively moving, and larval zebrafish have not been observed to swim backward. This is reminiscent of the collective behavior of bacteria in circular chambers where the fluid flow disturbed by the edge-swimming bacteria pushed back the middle-swimming ones (6). The interaction between water flow generated by the circling proportion of animals and the diameter and shape of the confinement is a point of interest which we will model and further test experimentally in the future. A better understanding of the mechanisms and interactions that give rise to the confined zebrafish collective motion will allow us to optimally direct their behavior by designing appropriate confining boundaries.

Here we share a new paradigm where collective motion can be induced by confinement in a model system amenable to genetic and neurobiological tools to investigate the underlying neural circuits. By understanding what influences this collective behavior and manipulating the enclosure scales and shapes in the future, we can determine fundamental interaction rules that would be widely applicable to other organisms and systems.

## Acknowledgements

We thank Dr. Simon Garnier and the Swarm lab for insightful suggestions and mentorship throughout the project, as well as Rafael Asfour from the Swarm lab for assistance implementing trackR. Mahathi Mohan Gowda built the behavior rig setup utilized for the experiments, along with members of the Severi lab and Dr. Christoph Gebhardt assisted in designing and building the behavior rig and gave input on analysis. We thank all members of the Severi lab past and present for excellent animal care and useful discussions throughout the project. Hassan Elsaid contributed through the NJIT Research @Home program. KES acknowledges support from NJIT startup funds. EL acknowledges support from the Simons Foundation as well as NJIT startup and seed funds.

## CONTRIBUTION TO THE FIELD

We provide the first observation of the transition from disordered to ordered collective motion of larval zebrafish by varying the number density in confined spacing. Studies of confined swimmer motion and interactions at the intermediate Reynolds remain sparse, and here we show that collective motion can be controlled by introducing surfaces and varying the number density. The study opens new research directions in many fields from active living matter to zoology and robotics. Understanding the interactions and emergence of collective motion of zebrafish in confinement at different densities may allow better experimental designs, with potential for some level of eventual control and direction of the behavior. In addition, this animal species lends itself to mechanical and neurobiological experimentation in the lab that can enable us to explore various mechanisms underlying collective behavior that may not be as accessible with other animals.

## SUPPLEMENTAL MATERIALS

**Supplemental Table 1:** Components for custom-built behavioral recording setup.

**SUPPLEMENTAL MATERIALS****Supplemental Table 1: Components for custom-built behavioral recording setup.**
| *Item* | *Model #* |
| --- | --- |
| Computer | Dell Precision 5820 Tower |
| High-Speed Camera | Mikrotron EoSens CL MC1362 |
| Lens | Fujinon HF35SA-1 2/3" 35mm F1.4 Manual Iris C-Mount Lens |
| Frame Grabber Drop-In Card | National Instruments PCIe-1433 |
| Bandpass filter | Midwest Optical BP850-49 from Graftek Imaging |
| IR LEDs | Infrared 850 nm IR LED Strip Light flexible strips connected with PN 3071 solderless connectors, all from Waveform Lighting |
| Cold mirror | Knight Optical 700FCQ160100 |
| Aluminum breadboard | Thorlabs MB4575/M |
| Rail | Thorlabs XT95-500 - 95 mm Construction Rail |
| Rail Baseplate | Thorlabs XT95P3 |
| Rail Mount | Thorlabs XT95P12/M |

**Supplemental Movie 1.** Density series from 5 to 130 larvae movie samples demonstrating representative behavior. Corresponds to Figure 2.

## Notes

### Competing Interest Statement

The authors have declared no competing interest.

